# Coupling abundance and spatial distribution for an unbiased and tractable beta diversity framework

**DOI:** 10.64898/2026.04.15.718733

**Authors:** Dingliang Xing

## Abstract

While fundamental for understanding community assembly, dominant beta diversity metrics suffer from severe sample-size dependence and lack tractable generative predictions. Here, we generalize Whittaker’s multiplicative index into a novel abundance-weighted framework. By proving this index strictly equals the expectation of the Good-Turing sample coverage deficit, we unify species abundance and spatial occupancy into a single, design-unbiased parameter. Furthermore, deriving exact neutral expectations provides process-based baselines to dissect beta diversity’s internal structure via additive decomposition into species (SCBD) and local (LCBD) contributions. Application to two large-scale forest plots empirically validates its strict sampling invariance and reveals non-neutral signatures. Crucially, our SCBD diagnostic reveals a striking ecological phase transition: rare-to-intermediate species are more spatially scattered than neutral drift predicts until reaching an aggregation onset threshold, beyond which dominant species exhibit supra-neutral aggregation, reflecting competition-colonization trade-offs. Concurrently, LCBD diagnostics successfully isolate the spatial footprints of environmental filtering from pervasive neutral noise. By avoiding the conflation of statistical estimation bias from genuine biological scaling and anchoring metrics in tractable models, this framework provides a rigorous toolkit that transforms our empirical understanding of spatial coexistence.

## 1. INTRODUCTION

A fundamental axiom of ecology is that biodiversity manifests along two intrinsically coupled dimensions: species abundance and spatial distribution (He & Legendre 2002; McGill 2010; Harte 2011). Although these dimensions are often analyzed separately—through species abundance distributions (SADs) or species occupancy distributions (SODs)—they are mechanistically inseparable. Beta diversity, broadly defined as variation in community composition across space (Whittaker 1960, 1972), emerges directly from this coupling, quantifying how local abundances scale up to generate regional patterns. Consequently, any rigorous measures of beta diversity must explicitly account for the joint effects of species commonness and spatial aggregation, while remaining invariant to the act of sampling itself.

The search for a unified framework to quantify this coupling, however, has historically been fraught with contention (Tuomisto 2010; Anderson *et al*. 2011). Following extensive debates on diversity partitioning (Lande 1996; Jost 2007; Tuomisto 2010; see Ellison 2010 for a forum), the field largely converged on the “numbers equivalent” framework based on Hill numbers (Hill 1973). This framework is algebraically elegant, not only because it unifies a continuum of diversity measures through the order *q*—which determines the sensitivity to rare versus common species—but also because it successfully bridges Whittaker’s two original and seemingly disparate conceptualizations of beta diversity (multiplicative beta and variance-based) under a unified mathematical formulation (Chao & Chiu 2016).

However, the hegemony of this framework has recently faced fundamental scrutiny. Critics argue that the mathematical axioms underpinning Hill numbers (e.g., the doubling property) often prioritize algebraic elegance over ecological realism (Ricotta & Feoli 2024), while the concept of “evenness” itself has been challenged as an unstable abstraction that fails to capture consistent biological signals (Alroy 2025). Beyond these conceptual concerns, the framework suffers from a critical practical limitation: Hill-number-based estimators do not, in general, provide design-unbiased estimates of beta diversity under finite sampling. Both empirical analyses and simulations demonstrate that these estimators can exhibit systematic dependence on sample size, such that observed variation in beta diversity may reflect differences in sampling intensity rather than true biological heterogeneity (Bennett & Gilbert 2016; Marion *et al*. 2017). Although rarefaction and coverage-based standardization are widely employed to mitigate these effects (Ricotta *et al*. 2019; Chao *et al*. 2023), they do not offer a general solution. Rarefaction curves frequently cross, leading to scale-dependent reversals in diversity rankings (Chase *et al*. 2018), and standardized metrics can remain biased by the underlying species abundance distribution (Xing & He 2019).

Null model approaches have therefore been widely adopted as an alternative strategy to control for sampling effects. A prominent example is the beta-deviation metric proposed by Kraft *et al*. (2011), which quantifies the standardized difference between observed beta diversity and a null expectation generated by randomizing species occurrences while preserving the SAD. While this approach effectively removes the sampling effect attributable to the SAD, it fundamentally changes the interpretation of the resulting metric. As argued by Xing & He (2021), beta-deviation measures the degree of departure from spatial randomness conditional on a given SAD, rather than beta diversity itself. By statistically controlling for abundance structure, null models isolate the spatial aggregation component but discard the contribution of abundance variation. This leads to a conceptual impasse: existing methods either conflate sampling noise with biological signal (as in Hill-number-based beta diversity) or artificially decouple abundance from spatial distribution (as in null-model-based approaches), whereas beta diversity arises from their intrinsic coupling.

Recent critiques have therefore advocated a return to pairwise dissimilarity, or equivalently a “raw variance”, perspective on beta diversity (Bennett & Gilbert 2016; Marion *et al*. 2017). Within this domain, the variance framework developed by Legendre & De Cáceres (2013) has emerged as a particularly influential approach. By treating the species-by-site matrix as a geometric object, this framework provides design-based, sample-size-invariant estimators of total beta diversity under its variance formulation. Moreover, it enables the additive decomposition of beta diversity into species contributions (SCBD) and local contributions (LCBD), which have become widely used diagnostic tools in biodiversity assessment and conservation planning (Heino & Grönroos 2017; Chen *et al*. 2026). However, to incorporate species abundances meaningfully, this framework relies on specific dissimilarity measures or data transformations (e.g., Bray-Curtis, Hellinger, or chord distances; Legendre & Borcard 2018). Although these transformations address issues such as double-zero symmetries and dominance scaling, they substantially increase mathematical complexity. Unlike simple incidence-based metrics, transformed abundance measures are difficult to derive analytically from first-principle ecological models.

This mathematical complexity points to a deeper limitation: the lack of theoretical tractability in abundance-based beta diversity. In macroecology, rigorous sampling theories have successfully linked species abundance distributions and spatial aggregation to beta diversity patterns, but these advances have been largely confined to incidence-based metrics (Plotkin & Muller-Landau 2002; Morlon *et al*. 2008; Xing & He 2021). Because incidence reduces abundance to detection probabilities, the expected values of these metrics can often be derived analytically from SAD models under explicit spatial assumptions. This tractability is largely lost for abundance-based metrics, with few notable exceptions (Condit *et al*. 2002). The challenge does not lie in modelling SADs perse, but in the mathematical structure of commonly used beta diversity metrics: whether based on Hill numbers (q > 0) or variance-based dissimilarities, the nonlinear aggregation of abundance frequencies renders analytical expectations under stochastic community models intractable in practice. As recently highlighted by Alroy (2025), standard abundance-based indices often fail to map consistently onto the parameters of underlying generative models, rendering them ineffective for parametric inference. As a result, the field faces a persistent dichotomy: incidence-based metrics admit rigorous probabilistic derivations but discard abundance information, whereas abundance-based metrics capture dominance structure but lack process-based theoretical expectations.

Here, we propose a framework that breaks this conceptual deadlock. We introduce a novel biodiversity parameter (denoted β for simplicity), which explicitly couples abundance and spatial distribution by quantifying the abundance-weighted mean probability of local absence. Using design-based inference, we show that the corresponding estimator is strictly unbiased with respect to sampling, regardless of sample size, the underlying species abundance distribution, or the degree of spatial aggregation. This formulation naturally admits additive decompositions into SCBD and LCBD within the variance framework. The SCBD component reveals our β as a generalization of Whittaker’s multiplicative beta, converging to proportional species turnover *sensu* Tuomisto (2010) when regional abundance is evenly distributed among species. The LCBD component establishes a direct link to Good–Turing sample coverage (Good 1953; Chao & Jost 2012), such that LCBD quantifies the coverage deficit of a local community relative to the regional species pool. Finally, we demonstrate that our β is theoretically tractable by deriving analytical expectations for the total index and its SCBD and LCBD components under the Unified Neutral Theory of Biodiversity (Hubbell 2001). Together, these results establish an abundance-based beta diversity framework that uniquely combines statistical unbiasedness with mechanistic tractability, enabling direct tests of macroecological processes against sampling-independent baselines.

## 2. THEORETICAL FRAMEWORK

### 2.1. Generalizing Whittaker’s Classic Index for an Unbiased Beta Diversity

We begin with the conceptual foundation of Whittaker’s classical multiplicative beta 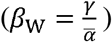, which remains the cornerstone of the field. Through monotonic transformation, this can be expressed as proportional species turnover (Tuomisto 2010): 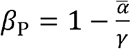. For a dataset of *S* species and *m* sites, this incidence-based index is mathematically equivalent to the mean probability of local absence across all species (Arita *et al*. 2008; Xu *et al*. 2015; Xing & He 2021):

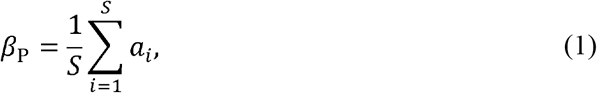

where *a*_*i*_ is the proportion of the *m* sites that is not occupied by species *i* (i.e., an estimator of the probability of absence). This metric treats all species equally, disregarding the fundamental dimension of abundance. More critically, this metric systematically depends on *m*, the sampling effort, and the bias is not trivially correctable (Bennett & Gilbert 2016; Marion *et al*. 2017; Xing & He 2019).

To explicitly couple abundance with distribution while rigorously resolving this sampling dependence, we generalize Eq. (1) by replacing the uniform weight (1/*S*) with the species’ regional relative abundance (*p*_*i*_). We define our target beta diversity parameter (*β*) as the expected abundance-weighted probability that a species is absent from a local community. Ecologically, this quantifies the likelihood that a specific individual, randomly sampled from the region, belongs to a species that is missing from a randomly selected local site. To estimate this parameter from a finite sample of *m* sites, we propose the following unbiased estimator, denoted as 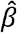 (or simply *β* in application):

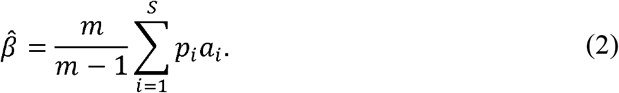

Here, *p*_*i*_ is the observed relative abundance of species *i* in the pooled regional sample, and *a*_*i*_ is the observed frequency of absence (the proportion of sampled sites where species *i* is not found). The index ranges from 0 (when all species are present in all sites, *a*_*i*_ ≡0) to 1 (when species are maximally segregated, i.e., each species is found in only one unique site, 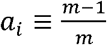. The term 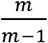 acts as a Bessel-like correction factor to ensure the estimator is design-unbiased. One can prove that the expectation of this estimator matches the true metacommunity parameter regardless of sample size *m*, spatial distribution, or species abundance distribution (i.e., 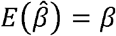, thus resolving the estimation bias typically associated with multiple-community beta metrics.

### 2.2. Decomposition and Connection to the Variance Framework

A key property of our linear additive formulation is that it allows for orthogonal decomposition into species- and site-specific contributions.

The contribution of the *i*-th species is defined as the term within the summation of Eq. 2:

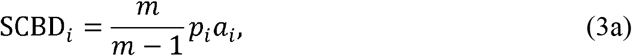

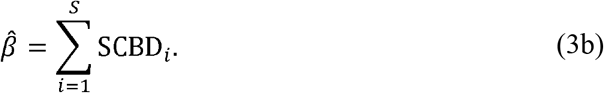

This formulation reveals a fundamental coupling between the species abundance (represented by relative abundance *p*_*i*_) and the spatial distribution (represented by local probability of absence *a*_*i*_). Conceptually, the relationship between SCBD_*i*_ and abundance *p*_*i*_ is expected to be unimodal. For rare species, SCBD_*i*_ is limited by low *p*_*i*_; for extremely abundant species, SCBD_*i*_ is limited by low *a*_*i*_ (as they occupy most sites). Consequently, beta diversity is predominantly driven by species of intermediate abundance that are common enough to be detected but spatially aggregated enough to generate turnover. However, we emphasize that this “hump-shaped” pattern is scale-dependent. In systems where the sampling grain is sufficiently small relative to the extent, even regionally dominant species may have non-negligible probabilities of absence (*a*_*i*_ ≫ 0), potentially leading to a monotonic increase of SCBD with abundance within the observed range. Formal theoretical derivation of the relation between SCBD and abundance is presented in the next section.

By rearranging the summation in Eq. 2 over sites *j =* 1, …, *m* rather than species, we derive the contribution of local site *j*:

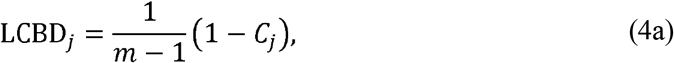

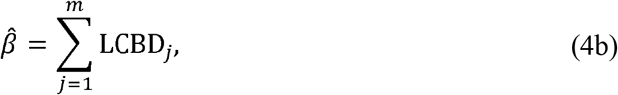

where 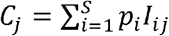 is the sample coverage of site *j* relative to the regional species pool, and *I*_*ij*_ is the indicator of presence. Thus, our framework establishes a direct theoretical link to Good-Turing frequency estimation (Good 1953): a site’s contribution to beta diversity is linearly proportional to its coverage deficit (1 - *C*_*j*_).

Ecologically, unique sites are those that fail to cover the regionally abundant species.

Critically, our estimator fully integrates with the coordinate-based variance framework proposed by Legendre & De Cáceres (2013). Our unbiased estimator 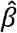 is mathematically equivalent to the average of all pairwise dissimilarities between distinct sites:

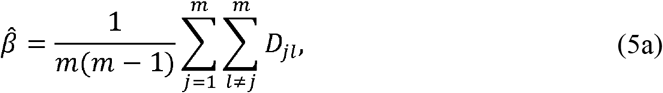

where *D*_*jl*_ represents the pairwise coverage deficit distance (PCD) between site *j* and site *l. D*_*jl*_ is derived directly from the general definition (Eq. 2) by setting *m* = 2:

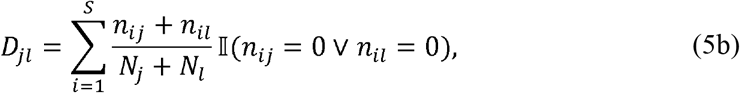

where *n*_*ij*_ is the abundance of species *i* in site *j, N*_*j*_ is the total abundance of site *j*, and 𝕀 (*n*_*ij*_ = 0 v *n*_*il*_ = 0) is an indicator function that equals 1 if species *i* is absent from one of the two sites (i.e., unique to the other) and 0 otherwise. Thus, *D*_*jl*_ quantifies the proportion of the pooled abundance of the pair that belongs to species unique to either site.

This metric offers a distinct perspective compared to other ecological distances utilized in the variance framework. Common coefficients such as the Bray-Curtis, Hellinger, or Chord distances quantify beta diversity by integrating differences in the abundance of all species, including fluctuations in the abundance of species shared by both sites. In contrast, PCD *D*_*jl*_ explicitly isolates the variation attributable to compositional uniqueness (coverage deficit). By strictly summing the abundance mass of species excluded from one site relative to the pair, it mathematically distinguishes the signal of species loss or replacement from the variation in abundance among shared species. This makes it a specific measure of abundance-weighted turnover driven by coverage deficits.

This equivalence (Eq. 5) provides a methodological advantage. Because 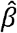 can be expressed as a pairwise dissimilarity matrix **D** = [*D*_*jl*_], it allows the rigorous application of standard multivariate statistical methods discussed by Legendre & De Cáceres (2013) directly to our abundance-coupled metric. These include coordinate-based ordination (e.g., PCoA) to visualize patterns of beta diversity, variation partitioning to quantify the relative roles of environmental and spatial predictors, and partitioning of beta diversity into within- and among-group components. The implementation of these analyses is statistically trivial using existing computational tools (e.g., functions cmdscale, dbrda, or adonis2 in the R package vegan; Oksanen et al. 2026). Due to space limitations, we do not expand on these specific multivariate applications here, but emphasize that our framework provides the theoretically unbiased distance matrix required to underpin them.

### 2.3. Theoretical Expectations and Sampling Theory

To provide a process-based baseline for inference, we derive the analytical expectations of *β*, SCBD, and LCBD. We employ a general framework combining a regional SAD with a spatial aggregation model.

#### The General Framework

Consider a metacommunity with *S* species constituting of *m* non-overlapping local communities of equal size. The expected value of our beta parameter can be expressed as the summation over the integer abundances of all species:

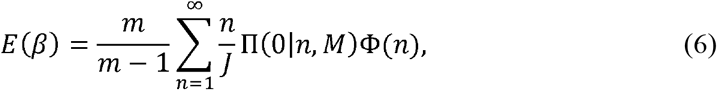

where Φ is the probability mass function of a species having regional abundance *n* (the SAD), Π (0|*n,m*) is the probability that a species with regional abundance *n* is absent from a randomly selected local community (the spatial distribution), and *J* is the total number of individuals in the metacommunity. While this summation is exact for discrete SADs, for continuous SADs in large systems it can be approximated by the integral 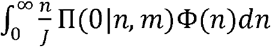, provided the SAD changes smoothly with *n*.

#### The Neutral Prediction

We instantiate this framework using the Unified Neutral Theory of Biodiversity (UNTB). Rather than assuming specific static distributions, we employ the predictions derived from the classic birth-death-immigration stochastic process (Kendall 1949; Hubbell 2001). The master equation predicts that the metacommunity SAD follows Fisher’s log-series distribution (Volkov *et al*. 2003), characterized by the parameter *x* (reflecting the regional effective birth/death ratio), where the probability of a species having *n* individuals is 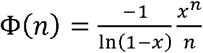. For each species *i*, the same stochastic model predicts that its abundance in a local community follows a negative binomial distribution (Kendall the 1949; Kitzes 2019), where the probability of local absence, conditional on regional abundance *n*, is 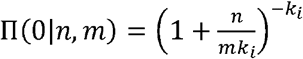. Under the neutral assumption, the species-specific aggregation parameter *k*_*i*_ can be expressed as a function of local birth/death ratio *x*′ and mean abundance 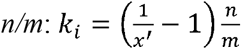. Substituting these process-based predictions into the general framework (Eq. 6), we derive the closed-form expectation for neutral beta diversity:

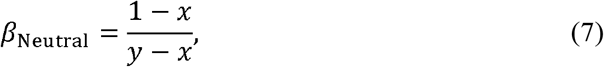

Where 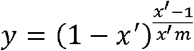. This formula demonstrates how speciation and neutral dispersal fundamentally shape the spatial structure of biodiversity. Under zero-sum dynamics, 1 − *x* is a measure of the per capita speciation rate (or speciation plus immigration from the external biogeographic pool for small-scale metacommunities). It is obvious from Eq. 7 that higher speciation results in larger beta diversity. Similarly, 1 − *x*′ quantifies the per capita local immigration rate; higher immigration connects local communities more tightly, resulting in smaller beta diversity.

The approximate variance of the neutral beta can be derived using the total variance formula (Barton & David 1959; Xing & He 2021):

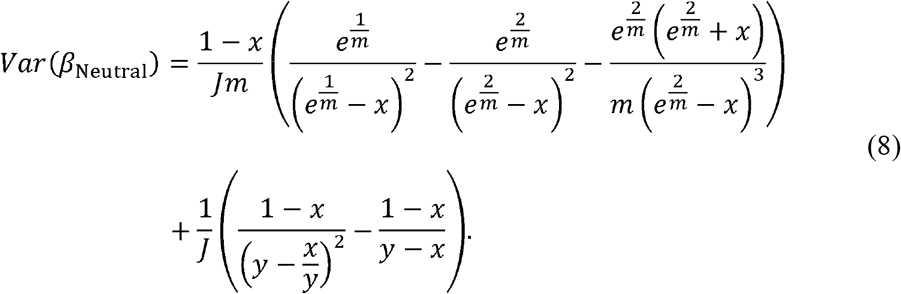

#### SCBD Under Neutral Theory

We further derived the explicit model for SCBD under neutral dynamics. Incorporating the NBD spatial structure, the expected contribution of a species with abundance *n* is:

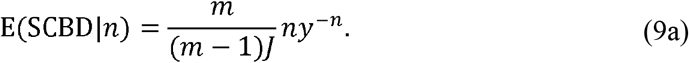

To facilitate hypothesis testing, we derived the sampling variance of SCBD. Since *n* (and thus relative abundance *p*_*i*_ = *n*/*J*) is considered fixed in this context, the variance of SCBD arises strictly from the stochasticity of spatial occupancy (i.e., the variance of *a*_*i*_) and the variance propagation from estimation of local parameter *x*′:

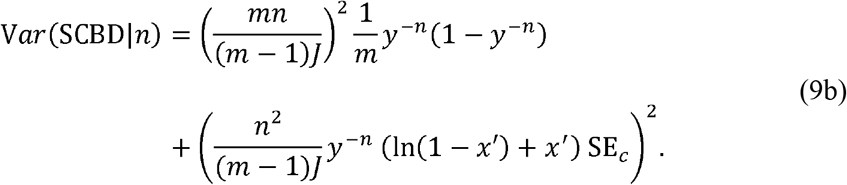

Here, SE_*c*_ is the standard error of parameter 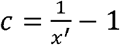, which can be estimated through fitting the linear relationship between aggregation parameter *k* and mean local abundance *μ*: *k* = *cμ* (see below). This analytical variance allows for the construction of confidence intervals around the theoretical SCBD curve, enabling the identification of species that deviate significantly from neutral expectations.

#### Theoretical Model for LCBD

The LCBD of a site (Eq. 4) represents the sum of relative abundances over the locally absent species. Under neutral theory, each species is independent (Haegeman & Etienne 2017), and thus the central limit theorem predicts that LCBD follows a Normal distribution 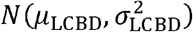 with:

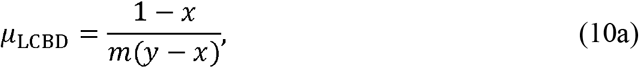

and

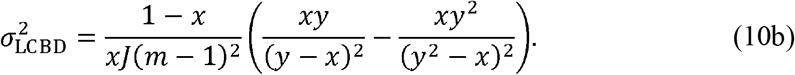

This Gaussian approximation adds to the toolkit a novel macroecological pattern for diagnosing the neutral model and identifying sites that are more unique than expected by stochastic dynamics alone.

#### Parameter Estimation in Practice

To apply these theoretical expectations to empirical data, the two key neutral parameters, *x* and *x*′, must be estimated from the observed community structure. The metacommunity parameter *x* is obtained by fitting Fisher’s log-series distribution to the regional SAD. The local parameter *x*′ is obtained by fitting the NBD model to species spatial distributions. Specifically, *x*′ can be estimated by simultaneously maximizing the NBD likelihood of observed local abundances, by incorporating the scaling relationship between the aggregation parameter *k* and the mean local abundance *µ* across species (*k* = *cμ*, where 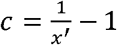).

## 3. EMPIRICAL APPLICATIONS

### 3.1. Data and Analyses

To demonstrate the statistical robustness and theoretical utility of our framework, we structured our empirical analysis to address two objectives: (1) validation of design-unbiasedness against sampling effort, and (2) diagnostic testing of neutral versus niche assembly mechanisms using our derived expectations and variance envelopes.

To achieve the first objective, we utilized census data from two large-scale forest dynamics plots: the 50-ha Barro Colorado Island (BCI) tropical forest in Panama (Condit *et al*. 2019) and the 35-ha Harvard Forest (HF) temperate forest in Massachusetts, USA (Orwig *et al*. 2022a). For BCI (Census 7, 2010), we analyzed 221,527 individuals with diameter at breast height (dbh) ≥ 1 cm across 300 species. For HF (Census 2019), we analyzed 52,763 individuals across 47 species. Each plot was divided into non-overlapping 20 × 20 m quadrats (*M*_*BCI*_ =1250,*M* _HF_ =875). Notably, the HF plot contains a contiguous swamp depression spanning 76 quadrats which were not included in the second census. Because our proposed occupancy-based metric mathematically accommodates empty local communities, we utilized all 875 quadrats to rigorously test for sample-size dependence. A random sample varying from 3 up to 400 quadrats was drawn without replacement 1,000 times to estimate total beta diversity (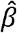, Eq. 2) and quantify estimation uncertainty. The design-unbiasedness of our proposed metric predicts that the sampling distribution will consistently center on the ‘true’ regional parameter calculated using all *M* quadrats. For comparison, we simultaneously calculated Whittaker’s classic beta index (Eq. 1), which is known to be sensitive to sample size.

To achieve the second objective, we generated a spatially explicit neutral landscape to serve as a strictly controlled theoretical baseline. We simulated a continuous-space neutral community using a backward-in-time coalescent algorithm implemented in the R package rcoalescence (Rosindell *et al*. 2008; Thompson *et al*. 2020). To ensure rigorous comparability, the simulation was parameterized to mirror the macroecological properties of the BCI plot. We populated a closed landscape grid with 650 × 325 individuals (totaling *J* = 211,250, approximating the BCI stem density). The fundamental biodiversity number was implicitly controlled by setting the per-capita speciation rate to *v* = 1.77 × 10 ^−4^ (calibrated to yield an expected regional richness of *S* ≈ 300). The local dispersal kernel width was set to *σ* = 26 grid units, corresponding to an equivalent distance of approximately 40 m in a 50-ha plot—a classic empirical estimate for dispersal limitation at BCI. This spatially explicit simulation successfully yielded a steady-state neutral metacommunity comprising 306 species.

For the diagnostic framework, the full empirical plots (BCI and HF) and the simulated neutral landscape were uniformly analyzed at the 20 × 20 m quadrat grain. Total beta diversity was additively decomposed into observed SCBD and LCBD vectors. Crucially, diagnostic testing against neutral expectations assumes a fundamentally habitable and homogeneous landscape; empty quadrats driven by severe abiotic constraints confound the effects of demographic drift with hard environmental filtering. Consequently, the 76 swamp quadrats in HF were strictly excluded from the mechanism-inference analyses, leaving *M* _HF_ =799 for the primary diagnostic models.

To construct the process-based neutral baseline, we estimated the two theoretical parameters from the observed community structure: the metacommunity parameter *x* was fitted from Fisher’s log-series distribution, and the spatial aggregation scaling parameter *c* (from the strictly neutral constraint *k*_*i*_ = *cμ*_*i*_) was estimated using composite marginal likelihood (CML). To handle the massive computational scale, CML was implemented via Template Model Builder (TMB). To rigorously account for the inflated degrees of freedom caused by abundance-dependent spatial autocorrelation—a pervasive artifact in spatial quadrat sampling— we eschewed analytical Hessian matrices and instead employed a spatial block Jackknife procedure to estimate a robust empirical standard error (SE_*c*_).

To avoid conceptual ambiguity, it is crucial to distinguish our process-based definitions of spatial structuring from classical textbook concepts based on Complete Spatial Randomness (CSR; Condit *et al*. 2000; Wiegand *et al*. 2025). Because real biological populations are always constrained by dispersal limitation, virtually all species appear highly aggregated when compared to CSR—especially rare species. In our framework, the reference baseline is the neutral steady state, which inherently assumes that spatial clumping is the natural consequence of limited dispersal and demographic drift. Deviations from this baseline yield a novel diagnostic scheme: Sub-neutral aggregation (negative SCBD residuals) indicates a species is less aggregated than a neutrally drifting species of identical abundance. Ecologically, this is the classic signature of inferior competitors, pioneer species, or those highly susceptible to conspecific negative density dependence. To avoid competitive exclusion or natural enemies, they must act as superior colonizers, broadcasting propagules widely and scattering individuals across a broader spatial extent than predicted by pure drift. Supra-neutral aggregation (positive SCBD residuals) characterizes dominant competitors or natural-enemy-insensitive species that successfully monopolize local patches, resulting in denser, more restrictive spatial clumps than neutral expectations.

To quantify this ecological phase transition across the abundance spectrum, we mapped the observed SCBD values against species abundances and employed a robust local polynomial regression (LOESS) to track the highly nonlinear residual trajectory. From this curve, we extracted two macroscopic metrics to quantify competitive spatial structuring: (1) The Aggregation Onset Threshold (AOT, *N*_cross_), defining the exact demographic tipping point where a species escapes sub-neutral aggregation and begins to exhibit niche-driven supra-neutral aggregation; and (2) The Niche Transition Slope (NTS), defined as the secant slope from the curve’s global minimum to the AOT, reflecting the acceleration rate of local dominance. Finally, to strictly evaluate the competition-colonization trade-off hypothesis, we applied a Right-Tail Spearman Rank Correlation Test to the species exceeding the minimum residual point, testing for the predicted monotonically increasing trend in aggregation.

Complementary to the species-level analysis, when diagnosing local spatial heterogeneity via LCBD frequency distributions, we decoupled the mean prediction from the variance shape. Specifically, we shifted the theoretical expectation curve (Eq. 10) to center on the observed total beta diversity. This “mean shift” ensures that our evaluation of local variance fluctuations is not confounded by minor systemic deviations in the overall mean prediction. Against this shifted baseline, each observed LCBD value was standardized into a Z-statistic using the theoretical neutral variance (Eq. 10b). Local quadrats demonstrating extreme compositional uniqueness—defined as those significantly exceeding neutral drift expectations (Z > 1.96)—were subsequently projected onto spatially explicit heatmaps. This procedure allows us to systematically pinpoint localized hotspots of non-neutral ecological turnover across the plot.

We developed an R package to realize all calculations, parameter estimations, and diagnostic visualizations (beta4s; https://github.com/dxinglab/beta4s).

### 3.2. Results

#### 3.2.1. Empirical Validation of Design-Unbiasedness

Our subsampling analysis empirically validated the strict design-unbiasedness of our generalized, abundance-weighted beta diversity metric 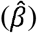. As expected, Whittaker’s classic incidence-based proportional species turnover (Eq. 1) exhibited severe sample-size dependence (Fig. 1a). Its estimates systematically underestimated the true regional beta diversity at small sample sizes; more critically, even at a massive sampling effort of 400 quadrats, a persistent deficit remained, demonstrating that its theoretical asymptote is practically elusive under realistic field conditions.

**Fig. 1.**
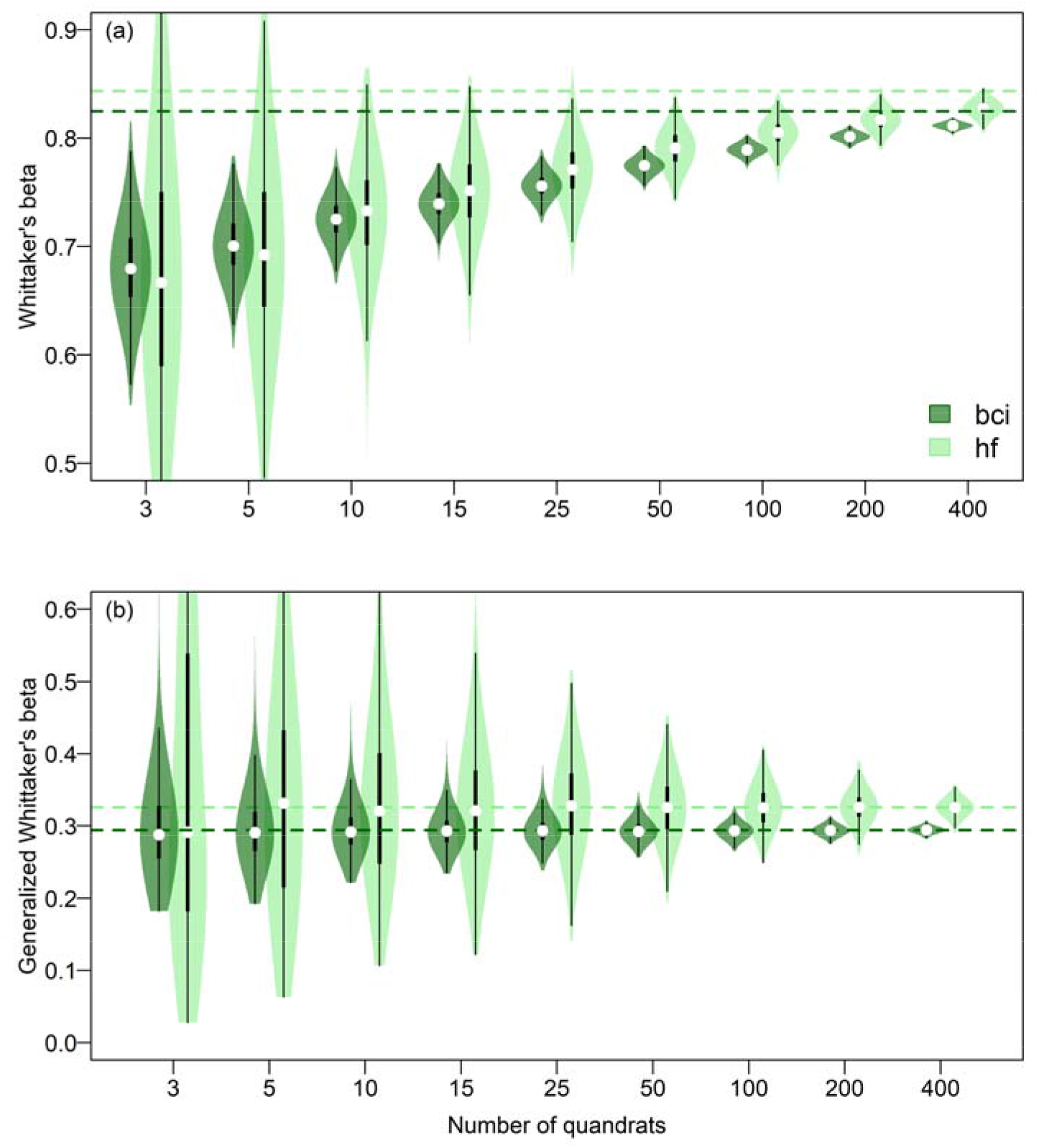
Violin plots showing the effect of sample size on estimates of different beta diversity measures. (a) Whittaker’s classical multiplicative beta diversity index, monotonically transformed to the so-called proportional species turnover (Eq. 1). (b) The new beta measure proposed in this study (Eq. 2). Dark and light green colors are for woody species from a 50-ha tropical rainforest in Barro Colorado Island, Panama (bci) and a 35-ha temperate forest in Harvard Forest, USA (hf), respectively. The dashed horizontal lines show the ‘true’ beta values calculated using the full data (i.e., the 1250 20×20 m quadrats for bci and 875 quadrats for hf). Each violin shows distribution of estimated beta values from 1000 repetitions of subsampling at a specific sample size (number of quadrats).

This systematic bias severely compromises comparative studies: due to differing accumulation rates, the apparent ranking of beta diversity between the tropical (BCI) and the temperate (HF) forests artificially reversed at lower sampling intensities.

In stark contrast, our proposed 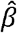 estimator (Eq. 2) remained perfectly invariant to sampling effort, completely eliminating this mathematical artifact (Fig. 1b). Regardless of the underlying community structure, the sampling distributions of 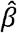 were consistently centered on the true metacommunity parameters (dashed horizontal lines). Notably, even under extreme undersampling scenarios (e.g., as few as 3 or 5 quadrats), the estimator yielded no systematic bias, with estimation precision naturally improving as sample size increased. Regarding this precision, we observed that the sampling variance was consistently larger for the HF plot. This elevated fluctuation reflects the strong spatial heterogeneity introduced by the contiguous swamp depression, which was deliberately retained in this validation step to demonstrate our estimator’s mathematical robustness to structural zeroes. Ultimately, this confirms that our framework successfully decouples the measurement of beta diversity from the confounding effects of sampling intensity.

#### 3.2.2. SCBD Diagnostics and the Ecological Phase Transition

When partitioning total beta diversity into species-specific contributions (SCBD), our process-based neutral baseline clearly demarcated niche-driven spatial structuring from stochastic demographic drift. For the simulated neutral community, the observed SCBD values and their standardized residuals closely hugged the theoretical expectations across the entire abundance spectrum, with the LOESS trajectory hovering firmly around the *Z* = 0 baseline (Fig. 2a, b).

**Fig. 2.**
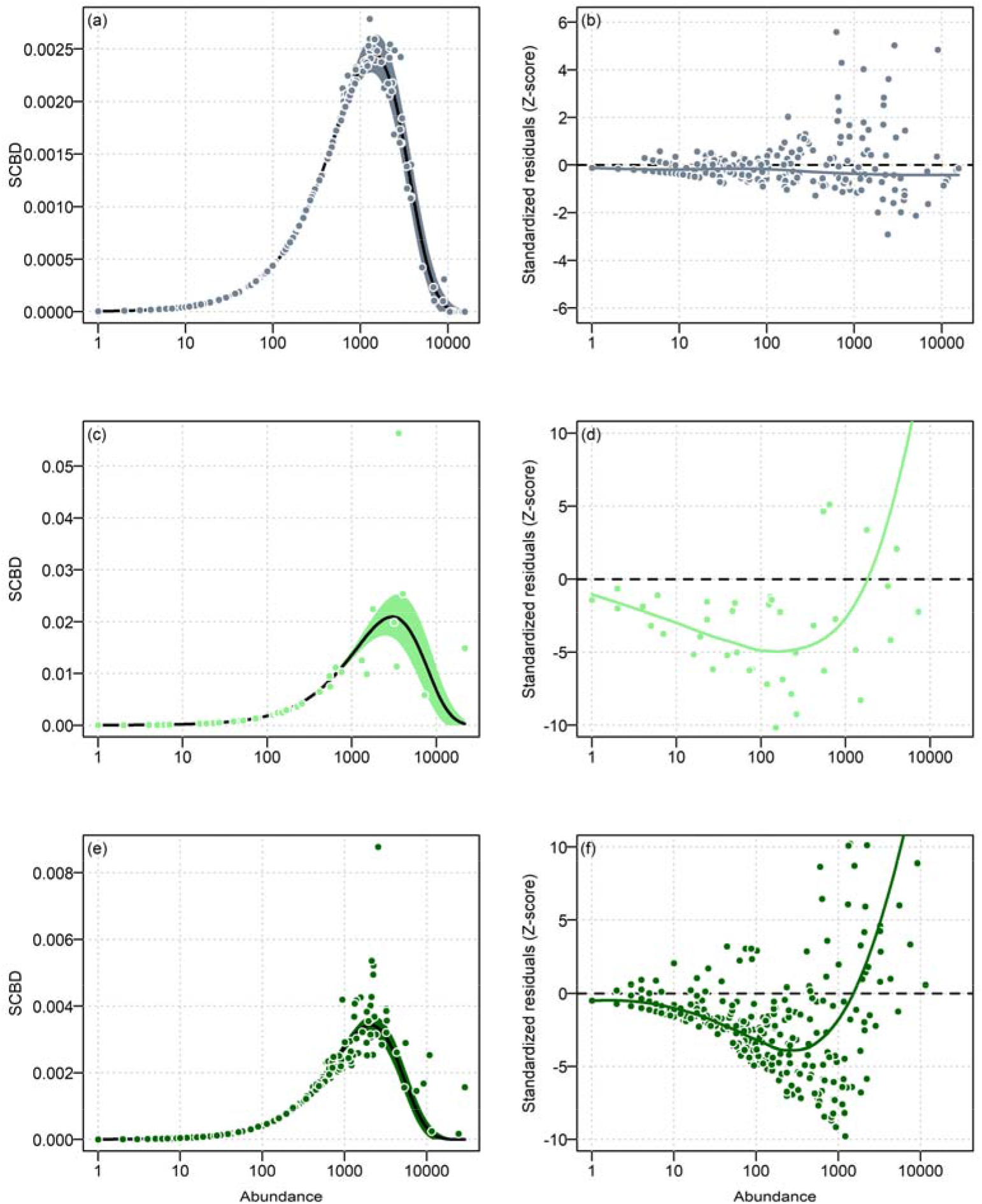
Species contributions to beta diversity (SCBD) and their standardized residuals across the abundance spectrum. (a, c, e) Observed SCBD against species abundance for a simulated neutral community (a), a 35-ha temperate forest in Harvard Forest, USA (c), and a 50-ha tropical rainforest in Barro Colorado Island, Panama (c). Solid black lines and shaded envelopes represent the theoretically derived neutral expectations (Eq. 9a) and their 95% confidence intervals (Eq. 9b). (b, d, f) The corresponding standardized residuals (*Z*-scores) of SCBD. The dashed horizontal lines indicate the strict neutral baseline (*Z* = 0), and the solid curves represent robust local polynomial regression (LOESS) fits tracking abundance-dependent structural deviations from the neutral expectation.

In the empirical forests, however, the standardized residuals revealed a striking, highly nonlinear “dip and rise” signature characteristic of strong niche assembly (Fig. 2c-f). At low to intermediate abundances, species consistently exhibited sub-neutral aggregation (negative *Z*-scores), indicating they are more spatially scattered than expected by neutral drift. This pattern aligns with the competition-colonization trade-off, where inferior competitors or density-dependent species act as colonizers to secure available micro-sites.

As abundance increased, the LOESS trajectory reached a global minimum before accelerating upward, crossing the neutral baseline at the Aggregation Onset Threshold (AOT, *N*_cross_). In the temperate HF plot, this ecological phase transition occurred at an abundance of approximately 1,833 individuals (Fig. 2d), while in the highly diverse BCI plot, the tipping point was observed at 1,529 individuals (Fig. 2f). Beyond this threshold, dominant species exhibited clear supra-neutral aggregation (positive *Z*-scores), forming stronger-than-expected spatial clumps indicative of local competitive superiority. The Niche Transition Slope (NTS), capturing the degree of departure from the neutral baseline, was 2.04 for HF and 2.23 for BCI. The Right-Tail Spearman Rank Correlation Test strictly confirmed this trade-off-driven spatial restructuring, revealing a highly significant monotonically increasing trend in aggregation for species exceeding the minimum residual point in both plots (HF: *P* = 0.013; BCI: *P* = 6.3×10^−7^).

#### 3.2.3. LCBD and the Detection of Extreme Spatial Uniqueness

At the local quadrat scale, the frequency distributions of LCBD provided a rigorous diagnostic for spatial heterogeneity, with a primary focus on the shape and variance of these distributions. For the simulated neutral landscape, the LCBD histogram closely followed our derived Gaussian approximation (Eq. 10; Fig. 3a). The proportion of quadrats significantly exceeding the upper threshold of the neutral expectation (*Z* > 1.96) was only 1.92%, aligning well with the expected Type I statistical error rate (e.g., *α* = 0.05). When projected onto a spatially explicit heatmap, these few high-LCBD quadrats were randomly scattered without forming coherent spatial clusters (Fig. 3b), perfectly reflecting the spatial footprint of pure demographic drift.

**Fig. 3.**
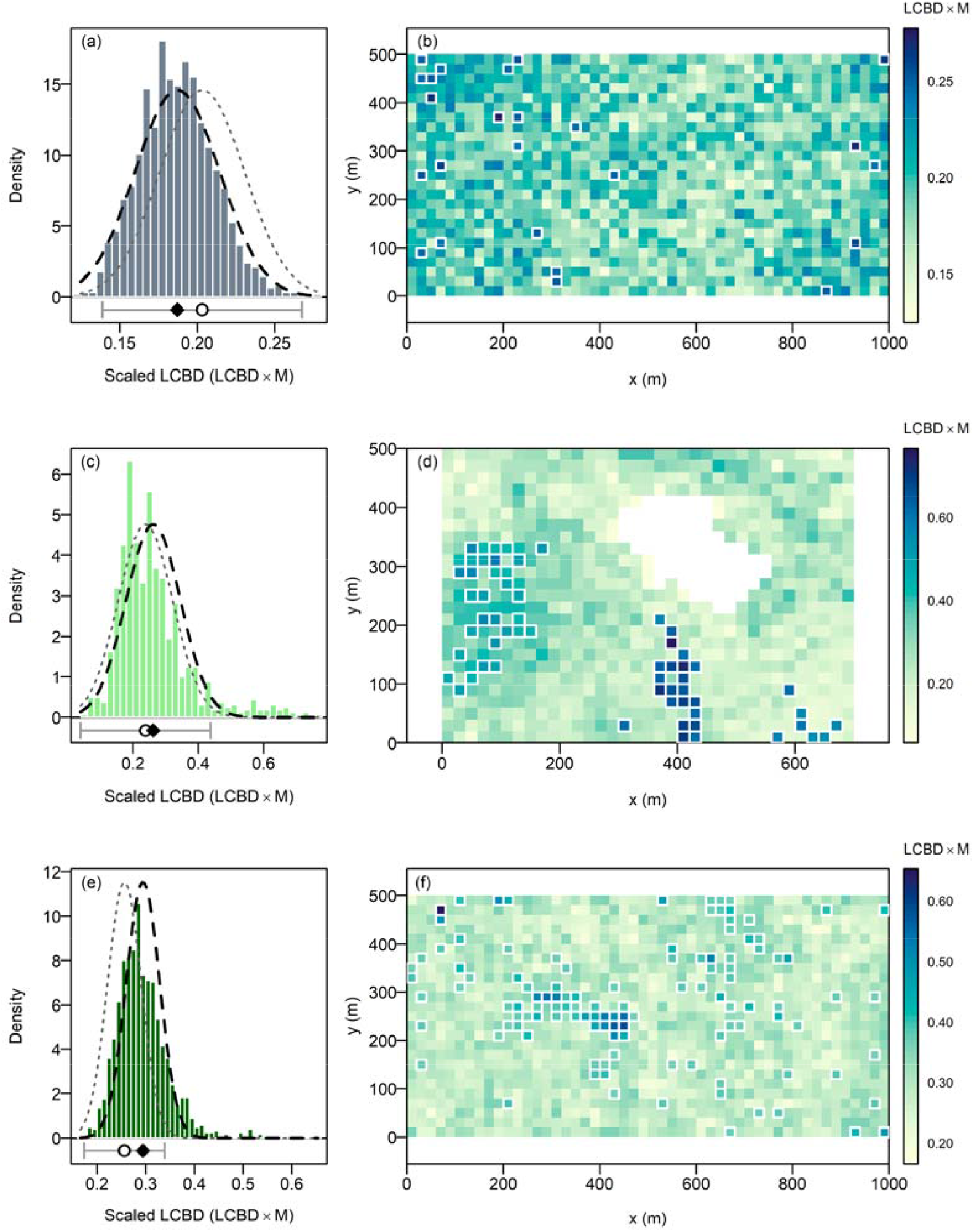
Frequency distributions and spatial maps of local contributions to beta diversity (LCBD). (a, c, e) Histograms of LCBD for a simulated neutral community (a), a 35-ha temperate forest in Harvard Forest, USA (c), and a 50-ha tropical rainforest in Barro Colorado Island, Panama (e). Dotted gray curves represent the parametric neutral expectations centered at the predicted total beta diversity (Eq. 7). Dashed black curves show the identical theoretical distribution shifted to center on the observed total beta diversity. At the bottom, circles and diamonds denote the predicted and observed total beta diversity, respectively. Horizontal error bars indicate ±1.96 SD derived from neutral sampling variance (Eq. 8). (b, d, f) Corresponding spatially explicit heat maps of LCBD. Darker colors indicate higher values. White bounding boxes highlight quadrats with significant departures from local neutral expectations.

For the empirical plots, although the total observed beta did not differ significantly from the neutral prediction, a clear qualitative departure in distribution shape emerged. Compared to the neutral expectation, both HF and BCI exhibited visually inflated variance and a pronounced right-skewed distribution, possessing a “long right tail” of highly unique quadrats (Fig. 3c, e). Specifically, the proportion of extreme non-neutral hotspots (*Z* > 1.96) reached 7.38% in HF and 9.28% in BCI, significantly exceeding the baseline stochastic expectation.

Crucially, these highly unique empirical quadrats were not randomly distributed. Spatial heatmaps revealed that they formed distinct, highly localized clusters (Fig. 3d, f). In HF, the most compositionally unique sites strongly aggregated in three specific sub-regions, likely reflecting the signature of land use history (Orwig *et al*. 2022b). Similarly, BCI displayed localized spatial blocks of high species turnover, with about half of the significantly unique quadrats coinciding with the center swamp habitat (Harms *et al*. 2001).

## 4. DISCUSSION

Our framework establishes a unified, design-unbiased, and theoretically tractable approach to abundance-based beta diversity. By explicitly grounding our metric in generative neutral processes, we transcend descriptive statistics to offer a rigorous diagnostic toolkit for identifying the precise footprints of deterministic community assembly.

### 4.1. Conceptual Unification: From Probability of Absence to Coverage Deficit

At its core, our proposed parameter (*β*) is both deeply rooted in ecological tradition and fundamentally novel. Conceptually, it is a direct generalization of Whittaker’s multiplicative beta, which, when expressed as proportional species turnover, essentially measures the unweighted mean probability of local absence across all species. However, by weighting this probability by regional relative abundance, our index seamlessly fuses the two inseparable dimensions of biodiversity: how common a species is, and how it is spatially distributed.

Furthermore, this formulation reveals an unexpected and profound equivalence with Good-Turing frequency estimation. While sample coverage has been widely popularized as a statistical tool to standardize diversity estimates via rarefaction and extrapolation (Chao & Jost 2012; Chao *et al*. 2023), our additive decomposition elevates coverage from a methodological corrector to an intrinsic macroecological property. The coverage deficit is not merely an artifact to be controlled for; it *is* the fundamental measure of abundance-weighted compositional turnover.

### 4.2. The Statistical Imperative of Design-Unbiasedness

A cornerstone of our framework is its strict design-unbiasedness. For statisticians, unbiasedness is the non-negotiable foundation of any robust inference, yet in community ecology, sample-size dependence has long been tolerated as an unavoidable nuisance. As Bennett & Gilbert (2016) and Marion *et al*. (2017) demonstrated, classical multi-community beta metrics are highly sensitive to sampling effort and the underlying gamma diversity. While it is sometimes assumed that these metrics will eventually plateau at a theoretical asymptote with sufficient sampling (Bennett & Gilbert 2016), our empirical subsampling (Fig. 1a) reveals that this asymptote is practically elusive even in heavily sampled large-scale forest plots.

Post-hoc dealing with naturally biased estimators is notoriously difficult. Rarefaction curves frequently cross, meaning that attempts to standardize biased beta metrics can lead to artificial reversals in diversity rankings depending on the chosen reference size (Chase *et al*. 2018). Recently, it has been proposed that the systematic decay of beta diversity estimators with increasing sample effort should itself be interpreted as a geometric ecological signal (Song *et al*. 2025). While we fully acknowledge the immense value of scale-explicit ecological analyses, we caution against conflating mathematical estimation bias with genuine biological scaling. True ecological scale-dependence—how beta diversity changes across physical grains and extents—can only be rigorously evaluated if the estimator itself is mathematically invariant to the arbitrary act of sampling intensity at a given scale. By providing an estimator that requires no rarefaction or coverage-based correction, our framework establishes the necessary unbiased baseline for comparative macroecology.

### 4.3. Beyond Statistical Attractors: A Multi-Dimensional Assembly Diagnostic

Tests of the Unified Neutral Theory have traditionally relied on a few macroscopic patterns (e.g., species abundance distribution, species area relationship, or distance-decay curve; Rosindell *et al*. 2011). Yet, ecology has increasingly recognized that these aggregate patterns often function as “statistical attractors” capable of being reproduced by mutually exclusive neutral and niche-based processes alike (Volkov *et al*. 2005; McGill *et al*. 2007; May *et al*. 2015). Our findings vividly corroborate this: the total observed beta diversity of our empirical forests did not significantly deviate from the neutral baseline, creating a false illusion of neutral assembly.

However, utilizing our additive decomposition (SCBD and LCBD) as a diagnostic “microscope” shattered this illusion. The variance approach to beta diversity introduced by Legendre & De Cáceres (2013) elegantly established the geometry of these local and species contributions, yet its ecological application has largely been restricted to post-hoc statistical regressions against environmental gradients or species traits. Our framework advances this legacy by providing analytically tractable expectations under a generative null model. This allows us to strictly quantify deviations from stochasticity before invoking external environmental data. As demonstrated in the empirical applications, focusing on the theoretical variance of LCBD allows us to definitively identify extreme non-neutral hotspots (Fig. 3). This effectively filters out the pervasive background noise of dispersal limitation, providing precise spatial targets for investigating deterministic environmental filtering or historical contingencies.

### 4.4. A Paradigm Shift in Spatial Aggregation

Perhaps the most transformative implication of our framework lies in the re-evaluation of spatial aggregation via SCBD diagnostics. Classical spatial ecology almost universally measures aggregation against Complete Spatial Randomness (CSR; Condit *et al*. 2000; Wiegand *et al*. 2025), a biologically implausible baseline under which virtually all species—especially rare ones—appear highly clustered due to inherent dispersal limitation.

By replacing CSR with a neutral steady-state baseline, we invert this traditional narrative. We found that low-to intermediate-abundance species consistently exhibit *sub-neutral aggregation*—they are significantly less clustered than demographic drift predicts. This provides powerful, scale-explicit evidence for the competition-colonization trade-off and local density dependence. As Lowe & McPeek (2014) articulated, dispersal is rarely a neutral lottery; it is an active trait mediating trade-offs. More recently, Kalyuzhny *et al*. (2023) demonstrated pervasive spatial repulsion among conspecific adult trees in tropical forests, driven by strong conspecific negative density dependence (CNDD). And quite a few studies suggested that rare species suffer stronger CNDD (Comita *et al*. 2010; Hülsmann *et al*. 2024). Our sub-neutral SCBD signature captures exactly this phenomenon at the community level: to survive predators, pathogens, or dominant competitors, vulnerable species must broadcast propagules widely, forcing a spatial distribution more uniformly scattered than expected by pure neutral drift.

Crucially, our robust LOESS tracking identifies a definitive “Aggregation Onset Threshold”. Beyond this demographic tipping point, CNDD is overridden, and dominant species achieve *supra-neutral aggregation*, actively monopolizing local patches. We anticipate that this phase transition threshold, alongside the Niche Transition Slope (NTS), will serve as novel, fundamental macroecological parameters. Future research must investigate how these parameters behave across different environmental regimes and, critically, how they scale with varying spatial grains, unlocking a more mechanistic understanding of how local coexistence scales to regional biodiversity.

## ACKNOWLEDGEMENTS

We thank Anne Chao for helpful comments on an earlier version of this work. DX was funded by the National Nature Science Foundation of China (32471623 and 32101280) and the Innovation Program of Shanghai Municipal Education Commission (2023ZKZD36).

